# *In vivo* quantitative high-throughput screening for drug discovery and comparative toxicology

**DOI:** 10.1101/2022.08.26.505462

**Authors:** Patricia K. Dranchak, Erin Oliphant, Bryan Queme, Laurence Lamy, Yuhong Wang, Ruili Huang, Menghang Xia, Dingyin Tao, James Inglese

**Affiliations:** National Center for Advancing Translational Sciences, National Institutes of Health, Rockville, Maryland 20850, USA; National Human Genome Research Institute, National Institutes of Health, Bethesda, Maryland 20817, USA

**Keywords:** drug discovery, infectious disease, genetic disorders, high-throughput screening, laser cytometry, model organisms, *C. elegans*, proteomics

## Abstract

Quantitative high-throughput screening (qHTS) evaluates the pharmacology of drug and investigational agent libraries for potential therapeutic uses, toxicological risk assessment, and increasingly for academic chemical tool discovery. Phenotypic HTS assays aim to interrogate molecular pathways and networks, often relying on cell culture systems, historically with less emphasis on multicellular organisms. *C. elegans* has served as a powerful eukaryotic model organism for human biology and disease by virtue of genetic conservation and experimental tractability. Here we describe a paradigm to enable *C. elegans* in qHTS using 384-well microtiter plate laser scanning cytometry. GFP-expressing organisms are used to reveal phenotype-modifying structure-activity relationships to guide subsequent life stages and proteomic analysis. *E. coli* bacterial ghosts, a non-replicating nutrient source, allow compound exposures over 7-days spanning two life cycles to mitigate complications from bacterial overgrowth. We demonstrate the method with a library composed of anti-infective agents, or molecules of general toxicological concern. Each was tested in 7-point titration to assess the feasibility of nematode-based *in vivo* qHTS, and examples of follow-up strategies were provided to study organism-based chemotype selectivity and subsequent network perturbations having a physiological impact. We anticipate a broader application of this qHTS-coupled proteomics approach will enable the analysis of *C. elegans* orthologous transgenic phenotypes of human pathologies to facilitate drug and probe profiling from high-impact chemical libraries for a range of therapeutic indications and study of potential toxicological signatures.

**Graphic Abstract:** 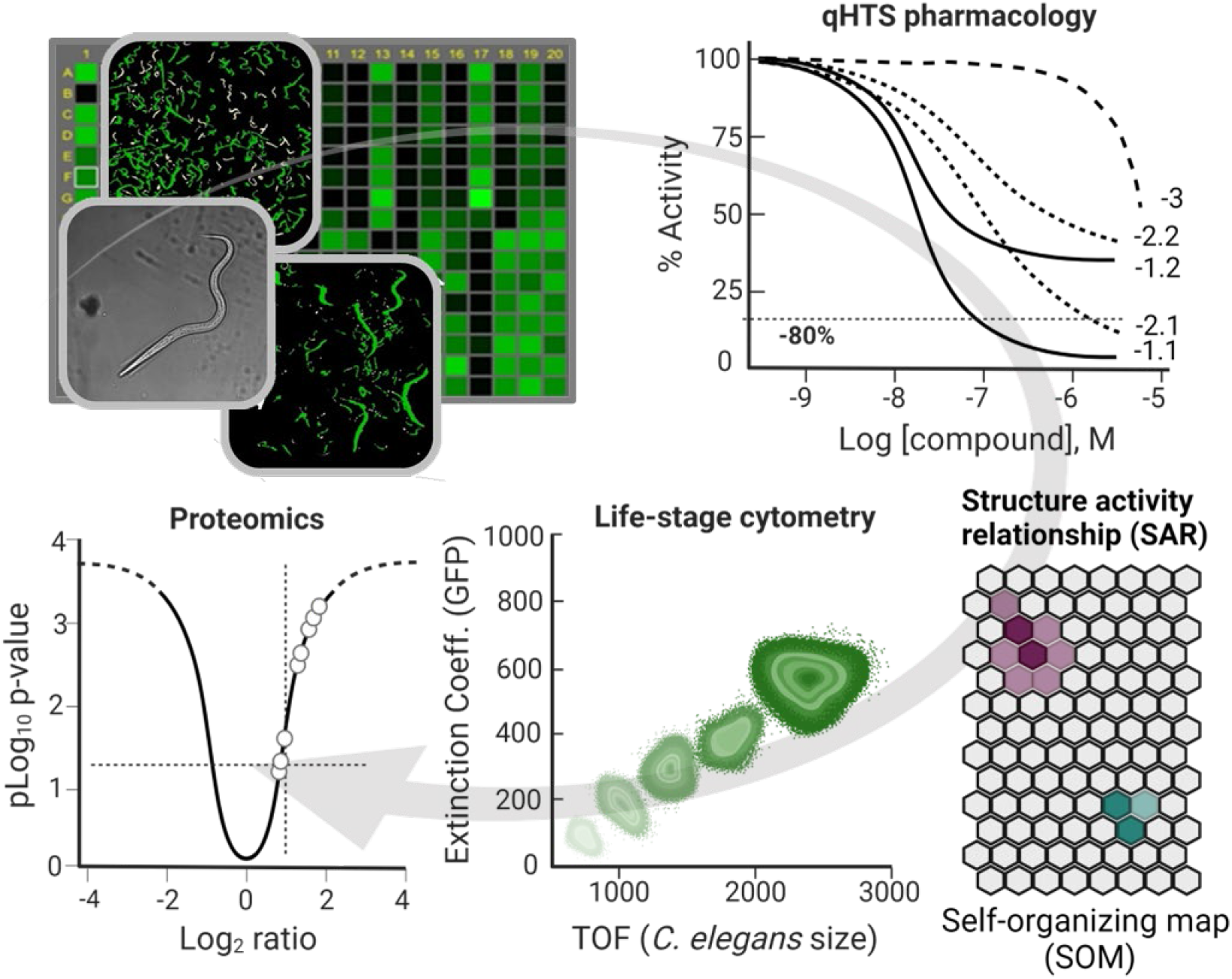

## Introduction

For over 40 years the nematode, *Caenorhabditis elegans*, has helped elucidate the molecular and genetic basis underlying a broad range of biological, developmental, and behavioral phenotypes (Brenner, 1974; Sulston and Horvitz, 1977). The organism’s short 3-day life cycle, minute size (1.5 mm long adults), anatomical transparency, ease of *in vitro* cultivation, and well-defined genetics are among its distinguishing experimental advantages (Riddle DL, 1997; Thompson et al., 2013). Bacteria serve as a live food source for *C. elegans*, and their rapid replication with *E. coli* permits the expansion and culturing of thousands of animals. As primarily self-fertilizing hermaphrodites that can cross with males, *C. elegans* furnish a highly controlled genetic environment. Along with a relatively small ∼20,470 protein-coding genome^1^, 35% of which have human homologs, the roundworm has proven to be a versatile model for human disease, including a means to interrogate rudimentary organ systems physiology (Kim et al., 2018). Disease studies in *C. elegans* have included those of the nervous system (neurodegenerative, neuromuscular, and axonal neuropathies), as well as metabolic diseases, protein misfolding and aggregation disorders among others (Caldwell et al., 2020; Kaletta and Hengartner, 2006; Kropp et al., 2021). *C. elegans* have also been used as a host organism for infectious microbes and for anthelmintic development against several parasitic worm species (Campbell, 1985; Ewbank and Zugasti, 2011; Sant’anna et al., 2013). Multiple studies have used *C. elegans* to interrogate the toxicology of potential anthelmintic agents based on viability, life span, and progression through various life stages. High-throughput screening (HTS) assays tracking these parameters to quantify compound efficacy and potency have been performed mainly using a single concentration treatment paradigm in a 96-well plate format (Breger et al., 2007; Petrascheck et al., 2009; Zhou et al., 2011).

In general, HTS has focused primarily on biochemical and cell-based assay formats, and efforts to incorporate the use of *C. elegans* have been comparatively moderate in scope and number of applications (Gosai et al., 2010; Inglese et al., 2007; O’Reilly et al., 2014). This, in part, can be attributable to growth condition incompatibility with large-scale screening, the amount of compound needed for dosing in traditional treatment paradigms, and labor-intensive handling of the worms (Brenner, 1974; Kwok et al., 2006). However, RNAi screening in *C. elegans* led to the progression of liquid culture in a 96-well microtiter plate format that was rapidly adapted to small-molecule drug screening (Lehner et al., 2006; Moy et al., 2006). The incorporation of automated liquid handling systems, image acquisition, and high-throughput data algorithms for image analysis and the miniaturization of assays to 384-well format have increased the efficiency of screening efforts in this whole organism system (Gosai et al., 2010).

A significant drawback in the implementation of HTS with *C. elegans* remains the potential for contamination or inconsistencies introduced from the bacterial food source over multiple day time courses. Bacterial overgrowth in wells due to metabolically impaired, stressed, or dead worms can impact movement and introduce environmental stress to the living worms or interfere with the data acquisition of specific phenotypes. Additionally, concern that effective compound concentrations may be reduced when using live bacteria due to metabolism or degradation may result in the need to use higher compound concentrations (O’Reilly et al., 2014).

Accordingly, heat-inactivated or antibiotic-treated *E. coli* have been employed to avoid these confounding effects (Gosai et al., 2010; Zheng et al., 2013). In an alternative approach, we utilized *E. coli* “ghosts” that can be prepared in bulk and stored for months to years to serve as a comparative food source for *C. elegans*. Bacterial ghosts (BGs) are cellular membrane envelopes from Gram-negative bacteria generated from the controlled expression of a recombinant PhiX174 lysis gene *E* initially employed for vaccine development and expanded as a vector for adjuvant, antigen, and drug delivery (Kudela et al., 2010). However, their use as a potential food source for *C. elegans* had not yet been reported.

The majority of HTS experiments involving *C. elegans* rely on microscopy-based high-content screening (HCS) to measure a phenotypic outcome, often by image analyses. While indeed a powerful measurement technology, HCS in this setting generally requires the acquisition of multiple fields to image a given well, accompanied by plate read times that can approach an hour or more. With wide-field imaging devices, a balance of speed and image quality is especially pertinent for fluorescent signals which must have sufficient detection intensity (Buchser et al., 2014). Exceptionally large data files from HCS can present a computationally intensive analysis effort, requiring complex algorithms to recognize and quantify the phenotype.

A sensitive and efficient option for HCS is microtiter plate-based laser scanning cytometry (LSC), a technology conceptionally comparable to flow cytometry (Zuck et al., 1999). In LSC, cells, particles, or more complex objects having intrinsic fluorescence above the background are detected and parametrically evaluated by the size, intensity and fluorescence distribution of select-gated objects (Kamentsky and Kamentsky, 1991). Independent of microtiter plate well density, LSC instrumentation can capture and quantify multiparameter data from individual particles on-the-fly, gate data to segregate subpopulations within each well, and acquire TIFF images, that can be further analyzed with HCS analysis software, in approximately 5 to 8 min (Auld et al., 2006). LSC is highly adaptable to whole organism screening when the entire body, an organelle, or a protein of interest is fluorescently labeled. Since its introduction, LSC has enabled a range of ligand binding and cell-based assays (Auld et al., 2006; Burgess et al., 2020; Clare et al., 2019; Sherman and Pyle, 2013; Zuck et al., 1999).

The generation of large-scale compound library pharmacology constitutes the conceptual basis of the quantitative high-throughput screening or qHTS paradigm (Inglese et al., 2006). When applied to model organism-based phenotypic assays, the approach is enabled at the level of *in vivo* physiology. In this context, qHTS can, for example, categorize physiologically relevant chemical classes for subsequent evaluation in chemo-proteomics-driven network analysis, or support their toxicological assessment for potential human or environmental liabilities. Here we demonstrate examples of multicellular organism qHTS using *C. elegans* to prioritize pharmacological responses from >600 annotated anti-infective molecules for subsequent bottom-up proteomics analysis (Burns et al., 2021), and in the survey of >800 environmental chemicals studied across *in vitro* assays in the Tox21 program (Xia et al., 2018). The platform is enabled by a novel non-replicating bacterial food source and the multiparametric detection sensitivity and rapid scanning speed of LSC to measure a fluorescent protein-encoded *C. elegans* phenotype (Hobert and Loria, 2006).

## Results

### *E. coli* BGs support *C. elegans* growth

To establish a stable non-replicating *C. elegans* nutrient reservoir suitable for a multi-day qHTS experiment, we investigated the use of *E. coli* bacterial ghosts (BGs). We compared whether worms would actively feed, display normal life stage progression, and reproduce when grown on BGs. Microscopic examination of BG-fed worms revealed similar numbers and mobility to a mixed population of worms grown on OP50 live bacteria. NGM plate-harvested worms analyzed by a COPAS FP-500 biosorter demonstrated an equivalent life-state distribution whether grown on live bacteria or BGs (**Figure 1A_C**). DiBAC4(3), a bis-oxonol membrane potential-sensitive dye selectively taken up by depolarized bacteria, demonstrated BGs filled the digestive tract of the *C. elegans* (**Figure 1D**). N2 worms were grown on OP50 live bacteria or TOP10 BGs with or without DiBAC4(3) dye for five days at an ambient temperature of ∼22°C spanning approximately 2 full life cycles. In only BG-fed worms was fluorescence detected by COPAS biosorter analysis (**Fig. S1**), where fluorescence increased in intensity from L1 through adult. These experiments indicate that *C. elegans* can feed on BGs and that this novel food source sustains normal worm development and brood size.

**Figure 1.**
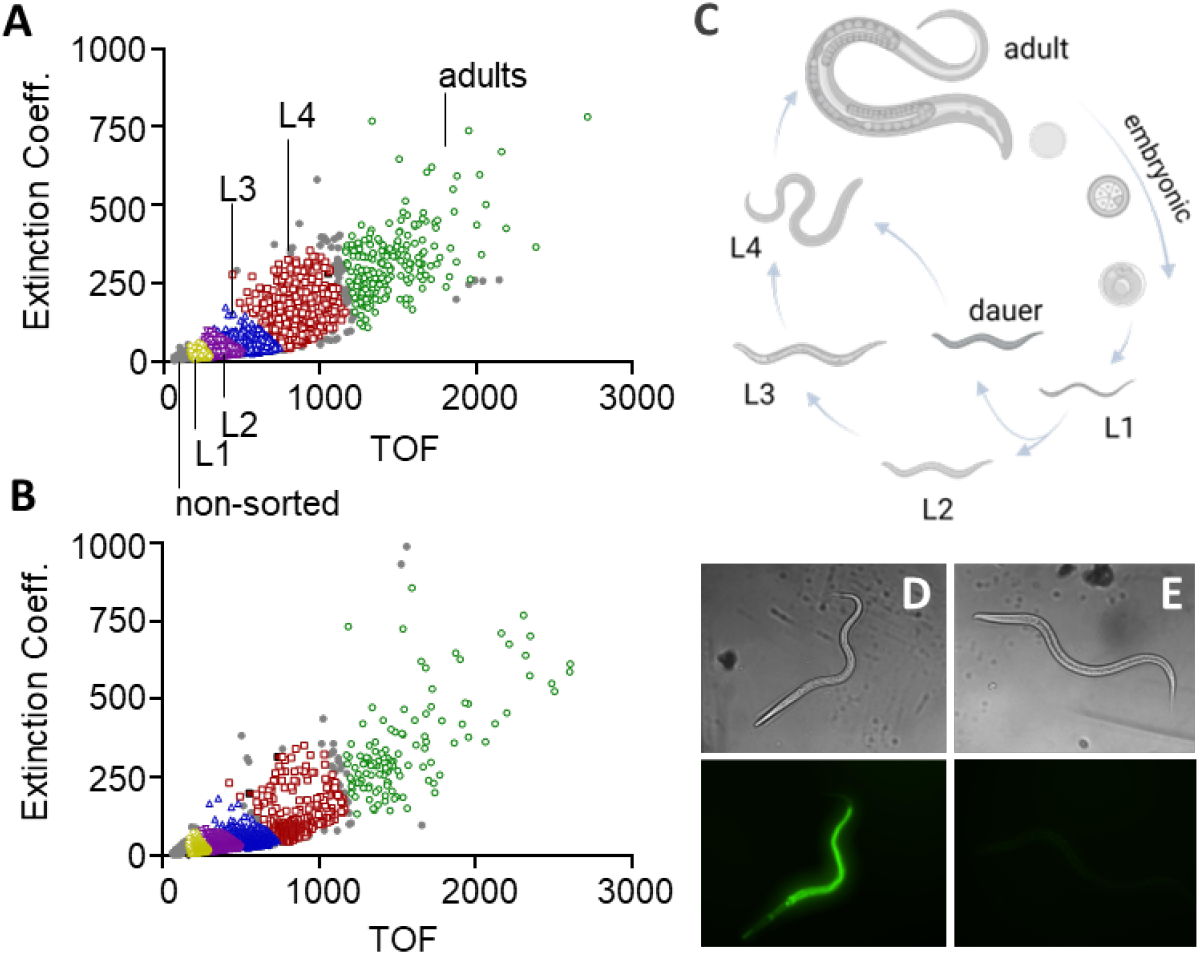
*C. elegans* life-stage analysis on a non-replicating bacterial nutrient. Scatter plots of strain N2 *C. elegans* life stage-sorted worms grown for five days on TOP10 bacterial ghosts (BGs) (**A**) or OP50 live bacteria (**B**). Life stages (**C**) were sorted on a COPAS biosorter based on Time-of-Flight (TOF) and Extinction Coefficient with various life stages shown as colored symbols and non-sorted objects shown as gray symbols. Representative brightfield and fluorescence microscopy images of L3 *C. elegans* strain N2 grown for five days on either TOP10 BGs (**D**) or OP50 live bacteria (**E**) pre-incubated with DiBAC4(3). Images were collected on an InCell 2200 imager with a 10x objective where brightfield (*top*) and FITC (*bottom)* filters were applied.

### 384-well laser scanning cytometry *C. elegans* viability assay

To verify BGs as a *C. elegans* growth media in 384-well microtiter plate format, the viability of various developmental stages of *C. elegans* strain PE254 that ubiquitously express GFP was assayed while grown on BG media or OP50 live bacteria with LSC. The viability of GFP expressing worms was quantified based on two parameters: (1) total GFP area (µm^2^) of gated objects, and (2) total gated GFP object number (“number of worms”) derived from integrated objects (**Figure 2A**), under either DMSO or 50 μM levamisole anthelmintic control treatment in 384-well format. Worms grown on live bacteria did not approach significance for HTS based on the Z-factor (Z’), a statistical measure to determine if the assay signal is reliably above background (Zhang et al., 1999), for any life stage or time point investigated. Worms grown on BGs demonstrated Z’ above the conventional HTS threshold of 0.5 for early larval stages, while later larval stages and adults had Z’ above 0.3 which supports qHTS feasibility (**Fig. S2** and **Table S1**).

**Figure 2.**
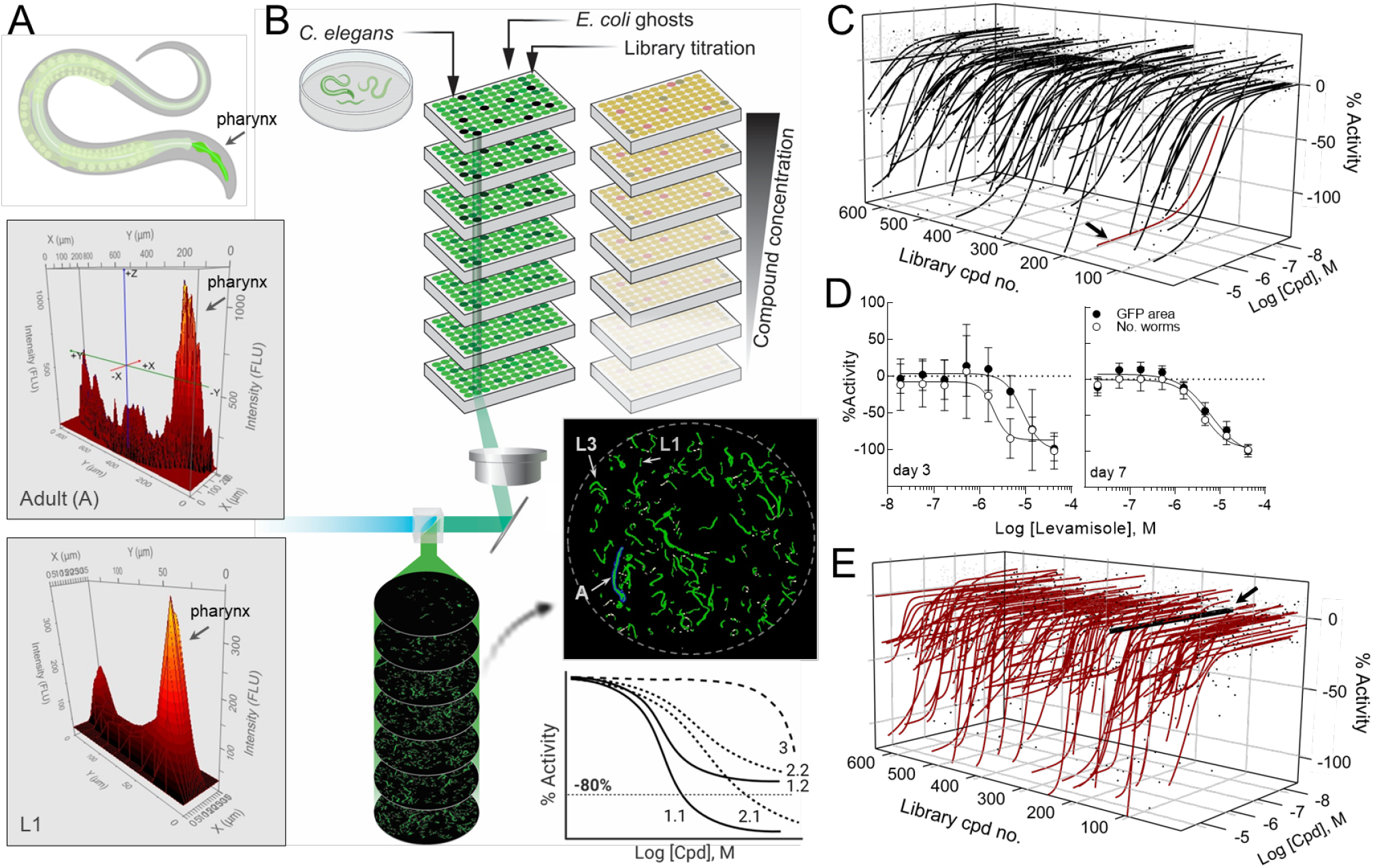
*C. elegans* laser-scanning cytometry qHTS. (**A**) Area (x-y) – intensity (z) plots for a representative adult, A and L1 *C. elegans* life stage, where the pharynx is clearly visible as the plot feature with the greatest GFP intensity. (**B**) Representation of the *C. elegans* qHTS process, highlighting the use of 384-well inter-plate compound titrations, laser-scanning cytometry output indicating the relative sizes of L1, L3, and adult worms, and general curve classification (CC) scheme for CC=1 (solid lines), CC=2 (dotted lines), and CC=3 (dash line) for dose-response curves (DRCs). (**C**) 3-axis plot of DRC profiles for GFP area of worms on day 7 for the 643 compounds in the NCATS Anti-Infectives collection; 4,501 dose-response values are displayed in blue dots. The 83 compounds with a decrease GFP area are plotted as black curves (**D**) DRCs for the levamisole screening control at day 3 (GFP area (•), 9.37 ±0.93 μM; No. worm (○), 5.96 ±2.40 μM) and day 7 (GFP area, 7.99 ±1.44 μM; No. worm., 5.74 ±0.91 μM). Error is SEM (n=21). Data was normalized to 41.7 μM levamisole as -100% activity and EC^50^ values were determined from logistic fits (GraphPad Prism) averaged from 6 replicates. (**E**) Compound library effect on HEK293 cell viability determined with Cell TiterGlo. Black arrows indicate the non-control ivermectin library sample DRC. Curves in 3-axis plot were fit using a four-parameter logistic regression software (see **Supplementary Methods** for details).

*C. elegans* were further assayed across a titration of three anthelmintic drugs: ivermectin, albendazole, or levamisole over 7 days. Each displayed differential life stage sensitivity to the drugs (**Fig. S3**). In general, worms grown on live bacteria exhibited more microtiter plate variability compared to those grown on BGs (**Figs. S2B** and **S3A, B**). Additionally, early larval stage worms appeared to progress through their life cycle and reproduce quicker on BGs compared to live bacteria. Nonetheless, accounting for growth rates, the relative pharmacotoxicity of the drugs was similar on worms whether grown on live bacteria or BGs (**Fig. S2C_F**).

Ivermectin, the most potent anthelmintic, displayed an ∼ 1 nM EC_50_ for all life stages grown on BGs by day 1 (**Table S2A**). Whereas albendazole, a microtubule poison that slows growth, and disrupts the larval cuticle and oocyte development has been reported to be toxic (EC_50_ >10 μM) on all life stages post 7-day treatment (Sant’anna et al., 2013), and this is consistent with our own observations for worms grown on live bacteria, though the drug was slightly more potent (EC_50_ = 500 nM to 4 μM by day 7 for all life stages), having minimal toxicity earlier in the time course. Whereas worms grown on BGs presented significant toxicity as early as day 1 for L4 and adult worms, and by day 2 for most life stages (**Table S2b**), correlating with the more rapid life cycle progression and egg laying observed on BGs. Levamisole by contrast gave similar EC_50_ values in the low micromolar range for all life stages grown on both media across the 7-day time course (**Table S2c**).

### *C. elegans* laser scanning cytometry qHTS

An overview of the phenotypic qHTS configuration utilizing *C. elegans* grown on an *E. coli* BG nutrient source is shown in **Figure 2B**. Here, a library of 643 agents having a variety of anti-infective mechanisms was prepared as a 7-point inter-plate titration series in 384-well assay plates, to which nutrients and *C. elegans* were subsequently added. *C. elegans* viability was assessed on worm number and total GFP area per well, respectively, on day 3 (one life cycle) and day 7 (>2 full life cycles) post-treatment. As expected, both parameters gave a greater signal-to-background and improved assay statistics on day 7 compared to day 3, although only the GFP area approached a significant Z-factor(Zhang et al., 1999) on day 3 (**Table 1**). EC_50_ values determined from both parametric outputs for levamisole, the intra-plate screening control, were similar on day 7 and correlated well with published values (**Figure 2D**) (Qian et al., 2008). Additionally, three anthelmintic controls (ivermectin, levamisole, and albendazole) were included as inter-plate controls on one assay plate and read every 24 h over the 7-day time course. EC_50_ values, while slightly higher than previously observed (**Table S2**), demonstrated activity at the expected time points (**Table 2** and **Fig. S3D**). In parallel, an HEK293 cell viability assay was conducted to estimate the compound toxicity window between mammalian cells and the nematode for all library compounds (**Figure 2E, Fig. S4** and **Table S3**) where ivermectin (black arrows, **Figure 2C** and **E**) illustrates an ideal species selective toxicity window.

**Table 1.**

| Output parameters | Assay parameters |  |  |  |
| --- | --- | --- | --- | --- |
|  | day 3 |  | day 7 |  |
|  | GFP area (FLU) | Worm number | GFP area (FLU) | Worm number |
| Signal : Background | 2.8 ± 0.4 | 2.2 ± 0.4 | 4.7 ± 1.1 | 5.2 ± 1.7 |
| Output signal | 50,400 ± 6,500 | 79 ± 16 | 90,600 ± 7,600 | 364 ± 26 |
| CV | 12.9 ± 3.0 | 19.8 ± 4.8 | 8.4 ± 1.7 | 7.1 ± 1.4 |
| Z' factor (median) | 0.29 ± 0.18 | -0.37 ± 0.81 | 0.57 ± 0.11 | 0.62 ± 0.13 |

Notes: Library size was 643 compounds tested across 7 concentrations. Experiment conducted in 21 inter-plate titration 384-well plates, plus two plates containing DMSO (0.42% final assay conc.) for field corrections. Data was analyzed against a control concentration of 41.7 μM Levamisole. Values represent Mean ± SD calculated from values based on N=10 DMSO and N=8 levamisole controls per plate.

**Table 2.**
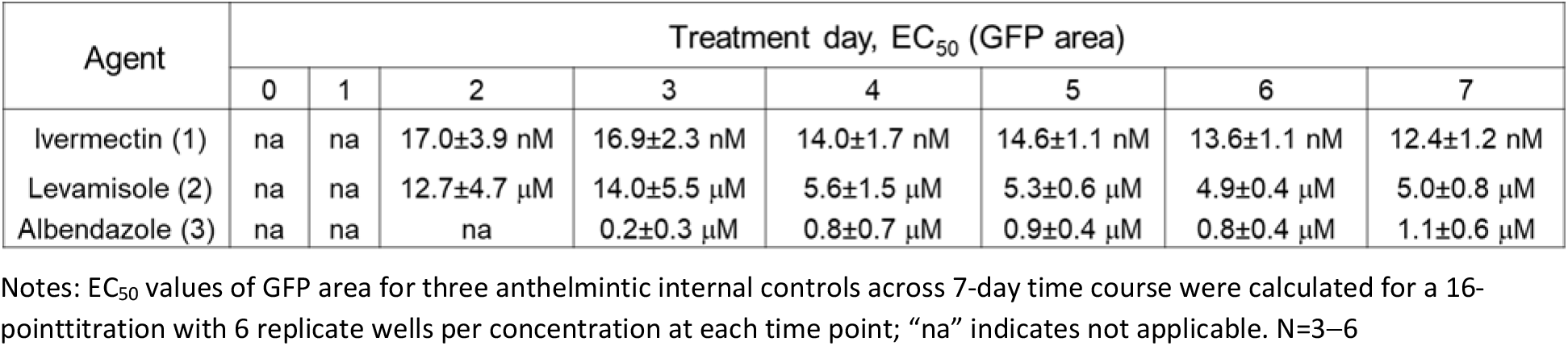

Analysis of GFP area on day 7 yielded 83 compounds that inhibited worm viability based on curve classifications of -1.1 to -3 with EC_50_ values ranging from 10 nM to >50 μM (**Table S3** and **S4**). Including ivermectin, two other anthelmintics, pyrvinium pamoate and L-tetramisole (levamisole), were also identified as active, whereas the fourth antiparasitic compound in the library, praziquantel, was not detected (CC=4). Active compounds were selected for follow-up analysis based on qHTS curve classification assignment (Inglese et al., 2006), ranking first for activity on days 3 and 7, followed by compounds with moderate day 3 GFP area activity, and having low to no toxicity in the human cell line counter qHTS.

### Anti-Infectives Follow-up Studies

Ninety-six compounds were selected for follow-up analysis based on the prioritization scheme above, 78 of which demonstrated activity on the parameter of GFP area on day 7 in the initial screen (**Table S4**). These compounds were tested across a 7-day time course in an 11-point intra-plate titration, analyzed by LSC, and worm number compared with high-content imaging (**Figure 3A**). Both LSC parameters, GFP area and worm number, were analyzed on day 7 with acceptable assay quality metrics, while the levamisole assay control demonstrated consistent EC_50_ values between the two measured parameters and those values calculated in the initial screen (**Fig. S5**). Of the 96 compounds analyzed, 73 reconfirmed activities on both GFP area and worm number at day 7, 19 were inactive and 4 compounds were selectively active with one parameter but not the other (**Table S5**). Over half the 96 molecules analyzed shared a common annotated MOA with at least one other compound in the follow-up subset.

**Figure 3.**
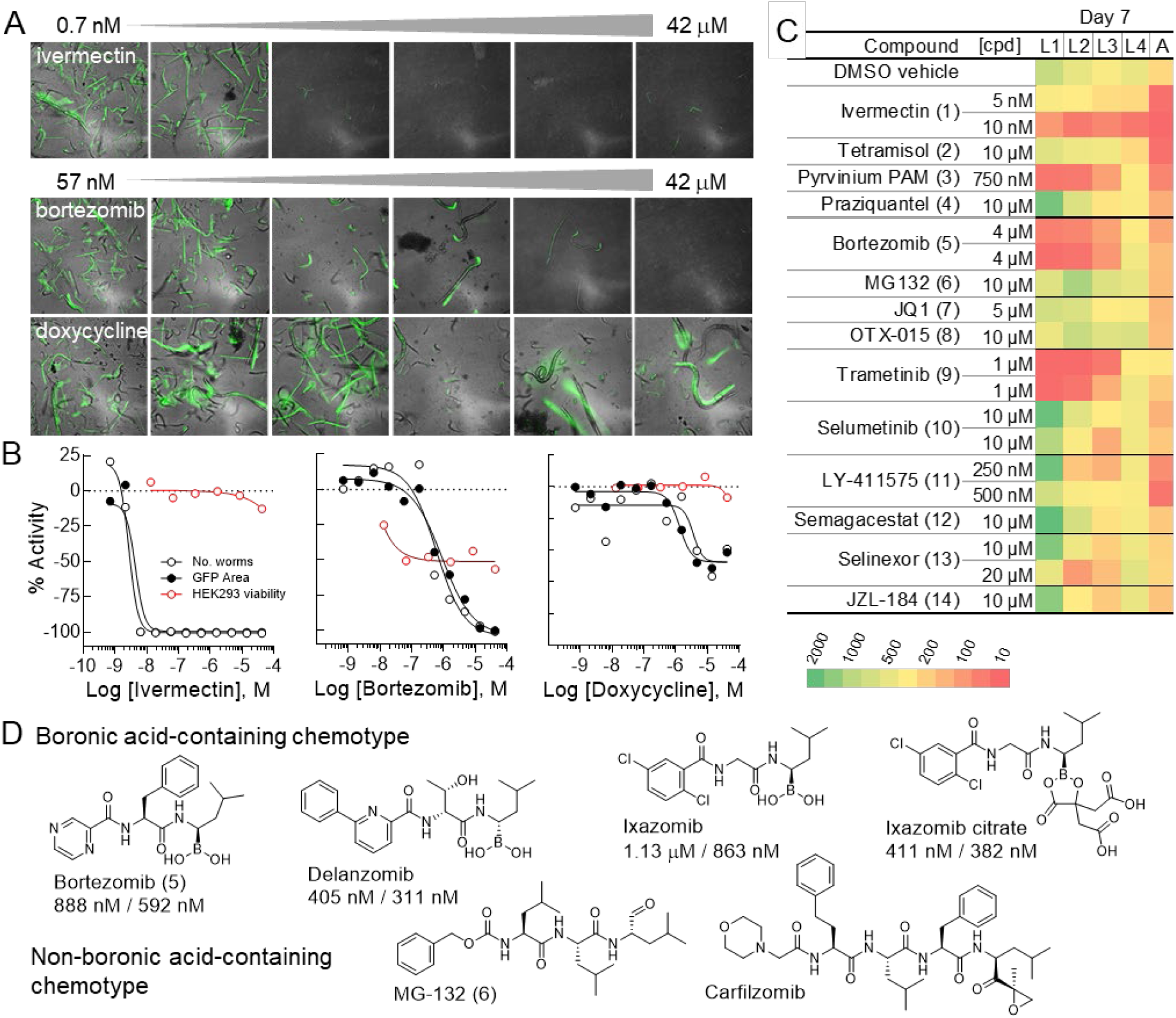
LSC-microscopy correlation and life-stage analysis. (**A**) Representative merged GFP and brightfield images obtained from the InCell 2200 across the active concentration range for the anthelmintic, ivermectin (*top*), the proteasome inhibitor, bortezomib (*middle*), and the antibiotic doxycycline (*bottom*). (**B**) Retest 11-point dose response curves for representative active molecules from three MOA classes identified in the primary qHTS and general cellular toxicity in mammalian cells (HEK293 cell line, ○). The two laser cytometer output parameters (GFP area, • and worm number, ○) measured on day 7 of the qHTS time course are plotted as data normalized to 41.7 μM levamisole screening control as -100% activity. Curves were fit in GraphPad Prism. (**C**) Day 7 life-stage distribution for larval stages L1-L4 and adult (A) as determined by COPAS flow cytometry. Worms were plated and treated as a mixed population of 2000 worms/well at day 0. The scale ranges from >15,000 (dark green) to <10 (red) animals. Compounds selected for evaluation on impact of life-stage viability: 1-4, antihelminth agents; 5-6, proteasome inhibitors; 7-8, bromodomain inhibitors; 9-10, MEK inhibitors; 11-12, γ-secretase inhibitors; 13, nuclear protein export inhibitor; 14, monoacylglycerol lipase inhibitor. (**D**) Boronic acid-based proteasome inhibitors and qHTS potency based on GFP area and worm number, respectively, and inactive non-boronic acid class of proteasome inhibitors represented in the library.

The three most active classes of molecules based on efficacy and potency included the anti-nematodal agents (3 of 3 identified), proteasome inhibitors (4 of 10 found active in the primary screen and all four reconfirmed activity), and bromodomain inhibitors (4 of 6 were identified as primary qHTS actives and reconfirmed activity). Additional compounds demonstrating effective cytotoxicity on *C. elegans* included an antifungal agent and one antimalarial agent, two MEK 1/2 inhibitors, both monoacylglycerol lipase inhibitors in the collection, and the two nuclear export inhibitors in the library (**Fig. S6A**). Also, five compounds listed as anticancer/antioxidant agents, two C-C chemokine receptor antagonists, and three DNA topoisomerase II inhibitors were found to have moderate activity. Among each of these classes were several inactive members suggesting species-related chemotype selectivity or potential off-target effects accounting for the observed activity (**Fig. S6B, Table S3**).

Two of the largest library categories included antibacterial/antimicrobial and antivirals. Several molecules from both were selected for follow-up where differential activity was observed within the respective MOA class plus the inclusion of several inactive molecules (**Tables S3** and **S4**). Two proteolytic secretase inhibitors demonstrated selective activity between gamma and beta-specific chemotypes, while deubiquitinase inhibitors and apoptosis activators were two other classes with variable activity in the *C. elegans* assay (**Fig. S6C**). Finally, 41 compounds had individual annotated MOAs within the follow-up subset, whereas tyrphostin A9, auranofin and oligomycin A had the best activity (**Fig. S6D**).

### Microscopy – laser cytometry correspondence

To confirm the correlation between LSC-derived DRCs and microscopy-based analysis, follow-up plates were imaged on the InCell 2200 on day 7 (**Figure 3A**). For each molecule, representative images from the active concentration range were visually inspected for worm number and health based on GFP and brightfield views. The potency of the various molecules appears to be highly correlated between the laser scanning cytometry- and microscopy-based imaging technologies, where efficacies based on the LSC DRCs across classes of MOA were also confirmed in the high content imaging, as visualized by the differential survival of worms under specific compound treatments (**Figure 3A** and **B**). Additionally, though the worms appear to be overlapping and clustered in the high content images, which can make segmentation difficult with HCS algorithms, the number of worms calculated on the fly in LSC with the optimized gating parameters accurately enumerated worms in each well which was very similar to the curves plotted from total GFP worm area (**Fig. S5**).

### Life stage flow cytometry analysis

For six mechanistic classes, we analyzed representative chemotypes (**Fig. S7**) at their ∼EC_75-80_ for effect on life stage viability after a 7-day treatment using the COPAS biosorter to enumerate organism subpopulations, summarized in the **Figure 3C** heatmap. Among the four re-tested anti-nematodal agents, differences in larval and adult populations were observed. We presume the dramatic treatment effects at 5 vs. 10 nM for ivermectin reflect the very steep dose-response of this drug as mirrored in the follow-up DRC pharmacology (**Figure 3B**), whereas the more graded hill slope observed (**Fig. S6A**) for tetramisol (cpd. 2) and pyrvinium pamoate (cpd. 3) permitted a more straightforward re-test concentration determination. As expected, the trematode parasitological agent, praziquantel (cpd. 4) had no effect on any *C. elegans* life stage, in agreement with the qHTS results (**Table S3**).

Clear species-dependent selectivity was observed among chemotypes (**Fig. S7**) targeting the same proteins or pathways. For example, as seen in **Figure 3C** the proteasome inhibitors, bortezomib (cpd. 5) vs MG123 (cpd. 6), and the MEK inhibitors trametinib (cpd. 9) vs. selumetinib (cpd. 10), either displayed potent larval-stage depletion or no discernable effect, respectively. Here, such chemotype-associated (e.g., **Figure 3D**) activity differences are not unexpected due to low sequence conservation between the human and nematode target proteins, though species-specific drug uptake and metabolism can also account for diminished or absence of compound activity.

### Label-free proteomics quantitation analysis

The annotated genome and pathway databases of *C. elegans* can inform the interpretation of phenotypic analysis pertaining to mechanism of action studies (Davis et al., 2022). To investigate this, we prepared digested peptides from protein extracted from ∼2,000 treated or control worms in biological triplicates, based on life stage analysis (**Figure 3C**), and subsequently the samples analyzed by HPLC-MS/MS (Burns et al., 2021). In brief, 2,645 protein groups and 37,992 peptides were identified, respectively, from 7 groups of samples **(Table S6**). Hierarchical clustering of the identified protein groups revealed clusters of up- and down-regulated proteins (**Figure 4A**), where molecules affecting life stage progression or number (cpds 5, 9, and 11) clearly separate from vehicle controls (DMSO or PBS) and inactive chemotypes (cpds 10 and 13) for targets of active compounds. Principle component analysis (PCA) from this data demonstrates a generally strong correlation of replicate treatments, especially for trametinib, bortezomib, and vehicle controls (**Figure 4B**).

**Figure 4.**
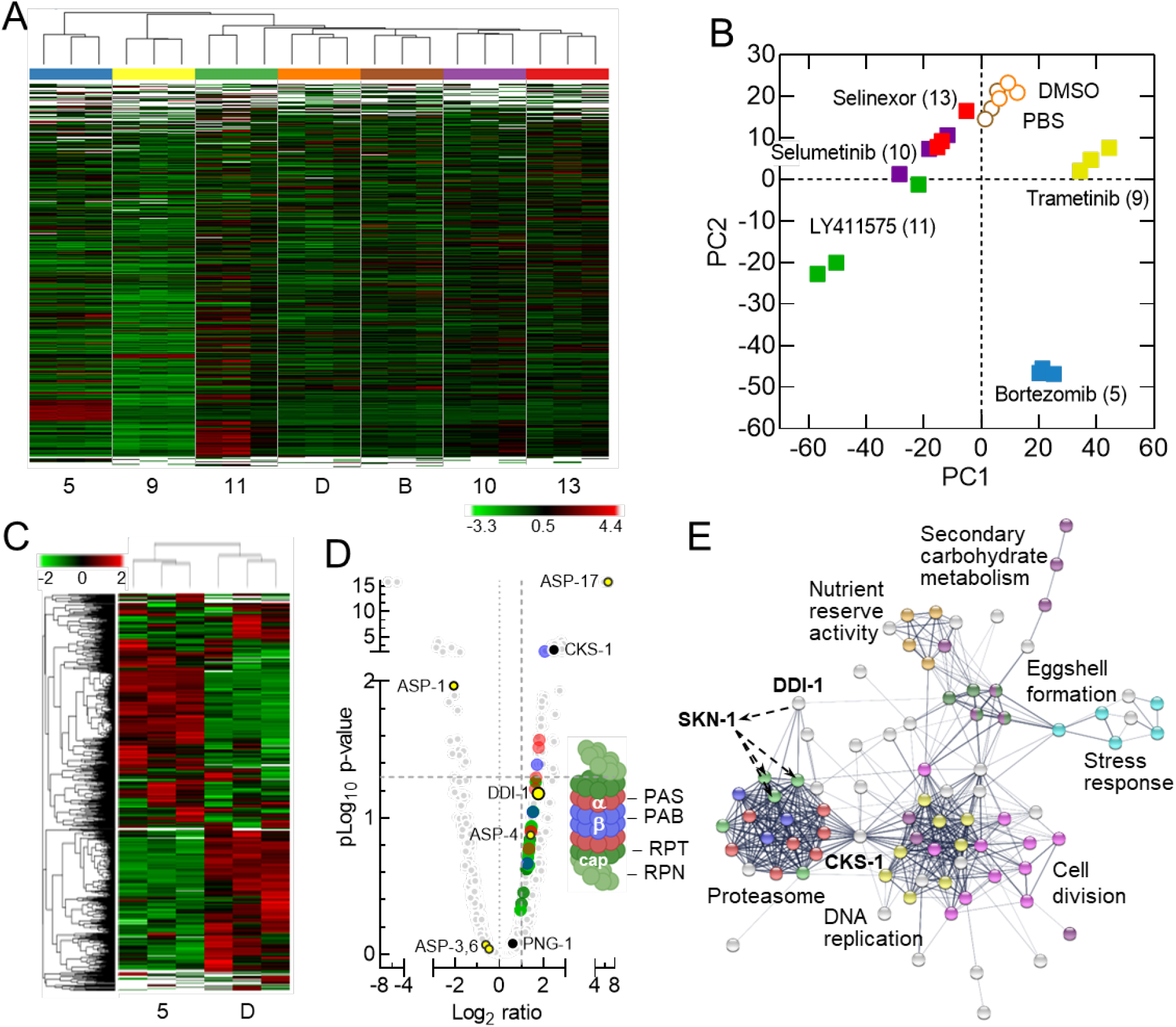
Proteomics analysis of drug-treated *C. elegans*. (**A**) Hierarchical cluster analysis using normalized protein abundance derived from a bottom-up quantitative mass spectroscopy-based proteomics analysis of *C. elegans* following a 7-day treatment with the indicated compounds or controls (D, DMSO or B, buffer). Label-free quantitation was applied for each group comparison with three biological replicates for each compound treatment and control condition. (**B**) Principal component analysis from 2,645 protein groups identified from 37,992 peptides for experimental triplicate determinations. (**C**) Hierarchical cluster analysis as in (a) for proteome changes induced by bortezomib vs DMSO vehicle control. (**D**) Volcano plot showing the statistical significance (pLog^10^ p-value) versus fold change (Log^2^ ratio) for 2,563 *C. elegans* proteins (light grey circles) after 7 days exposure to 4 μM bortezomib, 5, vs DMSO control. The 2-fold abundance increase (right of the vertical dashed line) are highlighted for proteins associated with the α, β and cap subunits (red, purple, and green circles, respectively) that comprise the proteasome (right inset). Open circles (○) are aspartyl proteases; Black circles (**•**) are proteasome-associated network proteins. The horizontal dashed line indicates the 0.05 p-value threshold. (**E**) A protein–protein interaction network constructed with STRING (*Search Tool for the Retrieval of Interacting Genes/Proteins*) ver. 11.5 using up-regulated proteins from the volcano plot (panel D). Coloring of the proteasome node correspond to the those used in panel D. The presence of skn-1 is inferred.

As an example of deeper phenotype interrogation, we examined the well-replicated bortezomib (cpd 5) induced proteome perturbations in finer detail. A direct comparative hierarchical clustering of the identified protein groups from bortezomib and DMSO treatments, allows the identification of proteins with statistically significant changes in abundance. From these proteomic changes, cellular consequence of proteasome inhibition was used to further encircle additional relevant processes (**Figure 4C** and **Table S7** and **S8**). These included, for proteins observed to increase in abundance by 2-fold or greater (**Figure 4D**), all 7 subunits comprising the α (pas) and 4 of 7 β (pbs) subunits of the 20S core particle, and 9 of the 15 cap subunit proteins (rpt and rpn) that make up the 19S regulatory particle of the 26S proteasome (Davy et al., 2001). A STRING network analysis focusing on up-regulated proteins clearly categorizes the proteasome and additional causally linked cellular processes, particularly DNA replication and cell division through the significantly up-regulated cyclin-dependent protein kinase regulatory subunit CDC28 ortholog, CKS-1 (**Figure 4E**) (Szklarczyk et al., 2019). Several aspartyl proteases are observed to be up-regulated as well. Among these, DDI-1 is responsible for the *N*-terminal processing of the transcription factor SKN-1, required for proteasome gene expression (Kropp et al., 2021). While SKN-1 was not detected, but rather inferred here, PNG-1, critical to the *N*-glycosyl asparagine editing of SKN-1 generating the aspartyl SNK-1 precursor DDI-1 substrate was detected in the *C. elegans* proteome (**Figure 4D, Table S7** and **S8**).

The translational up-regulation and/or half-life stability of drug-targeted proteins or complexes, and associated networks demonstrate the organism’s complex drug-mediated loss-of-function response. The subtle perturbations comprising associated processes are revealed and illustrated from a STRING network analysis exemplifying means to probe the physiological basis of phenotype-altering chemical agents (Kale and Moore, 2012; Szklarczyk et al., 2019).

### *In vivo* qHTS toxicology

Finally, the pharmacological output of *C. elegans*-based qHTS has demonstrated use in a comparative toxicology setting in the evaluation of >800 environmental chemical agents with potential toxicity identified from the U.S. Toxicology in the 21st Century (Tox21) effort (**Table S9**). The qHTS performed with similar statistics as observed for the previous anti-infectives library screen (**Table 3**) and in this case, we used total GFP area to score the activity measured on days 3 and 7 post chemical exposure. These 887 compounds were clustered based on a structural similarity approximation using the self-organizing maps (SOM) algorithm (Kohonen, 2006), yielding 192 *k*-clusters comprising between 1 – 15 members (**Figure 5A** and **Table S10**). Of these clusters, 20 were significantly enriched with compounds that inhibited *C. elegans* growth or survival, with the ivermectin cluster k16.1 displaying the most potent response (**Fig. S8**). Additional instances of clusters displaying potent activity include *k*8.9 represented by DDT and its analogs, *𝓀*1.3 containing a range of pharmacologically active substances having a shared piperidinyl alkylphenone substructure, and the organophosphates, irreversibly acetylcholinesterase inhibitors (**Figure 5B_D**). This latter category of 20 phosphates, thiophosphates and phosphonothioate

**Table 3.**

| Output parameters | Assay parameters |  |
| --- | --- | --- |
|  | day 3 (GFP area) | day 7 (GFP area) |
| Signal : Background | 2.5 ±0.6 | 2.9 ±0.7 |
| Output signal | 59,000 ±9,700 | 88,000 ±8,700 |
| CV | 16.9 ±0.4.4 | 9.9 ±1.9 |
| Z' factor (median) | -0.19 ±0.44 | 0.25 ±0.18 |

Notes: Library size was 887 compounds tested across 7 concentrations. Experiment conducted in 18 intra-plate titration 384-well plates, plus one plates containing DMSO (0.42% final assay conc.) for field corrections. Data was analyzed against a control concentration of 41.7 μM Levamisole. Values represent Mean ± SD calculated from values based on N=16 DMSO and N=8 levamisole controls per plate.

**Figure 5.**
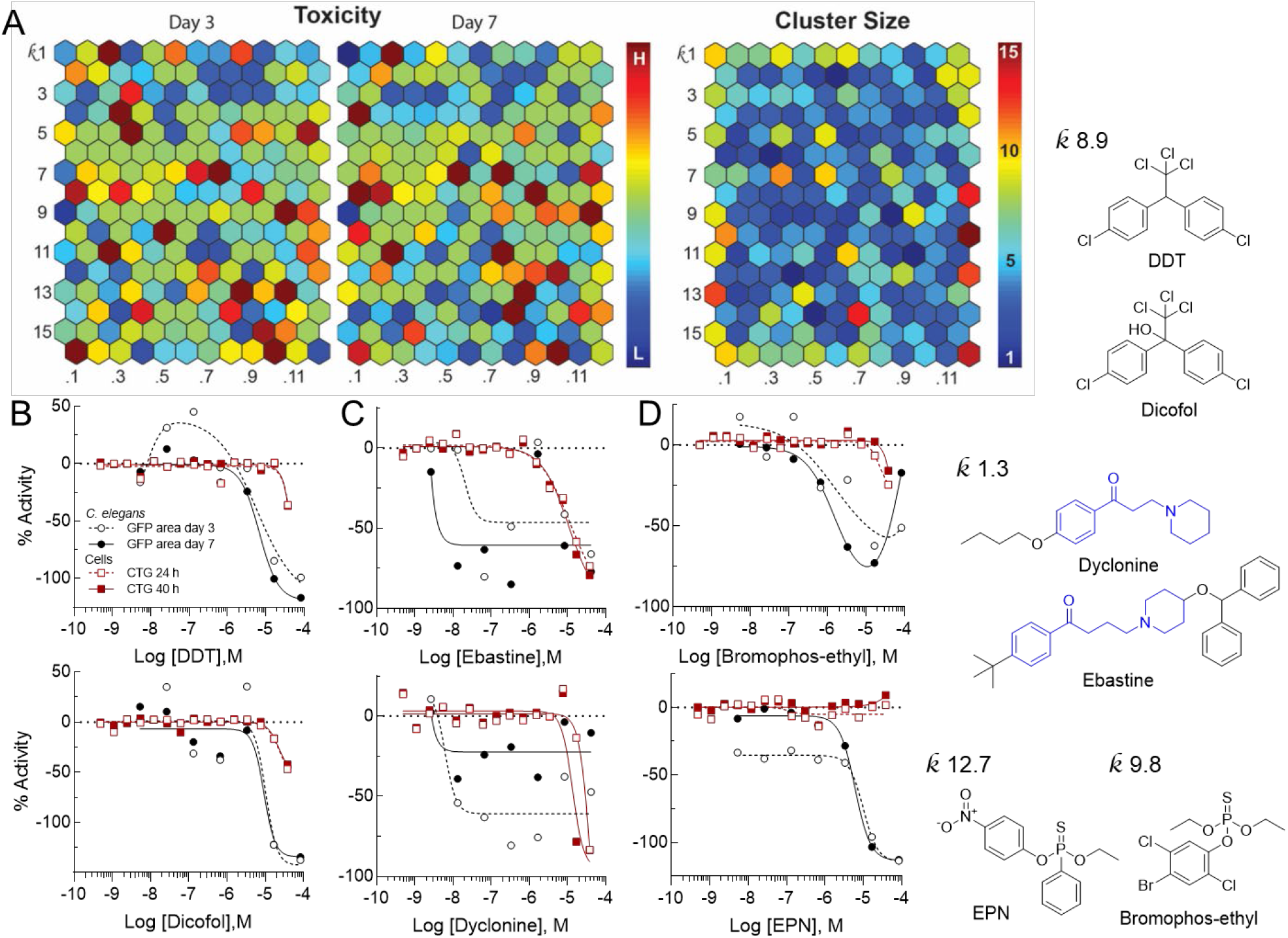
Analysis of Tox21 sub-library *C. elegans* qHTS output. (**A**) Structural – toxicity relationships as evaluated using the Self-Organizing Map (SOM) algorithm. In these heat maps, each hexagon represents a cluster of structurally similar compounds. Clusters enriched with potent active compounds (compared to the library average) are shown in red, and blue colored clusters are deficient of active compounds. Color scale values represent the log p value for each cluster that measures the statistical significance of enrichment from Student’s t-test. Representative dose-response curves plotted with the laser cytometer output parameter C. elegans GFP area (day 3, ○, and day 7, •) where data was normalized to 41.7 μM levamisole screening control as -100% activity, and for HEK293 cell viability data from the Tox21 program (24 h, ▫, and 40 h, ?) for several enriched clusters and chemotypes (**B**) *𝓀*8.9, endocrine disruptors, DDT and Dicofol; (**C**) *𝓀*1.3, piperidinyl alkylphenones (blue substructure), Dyclonine and Ebastine; and (**D**) organophosphates, EPN and Bromophos-ethyl distributed among different clusters.

EPN were distributed among 13 *𝓀*-clusters, illustrating a limitation of the SOM algorithm in chemotype identification. This is likely because the clustering algorithm, based on whole molecule similarity, does not prioritize substructure features such as a phosphate group when defining a cluster (**Fig. S9**). In addition to these compounds, 15 potential mitochondrial toxins, such as chlorfenapyr (*𝓀*7.8) and rotenone (*𝓀*12.2), were identified from a previous *C. elegans* study of the Tox21 program (Xia et al., 2018) and were also identified in the current *C. elegans* qHTS assay. As shown in **Table S11**, there is a very good correlation between *C. elegans* qHTS and previous ATP and larval growth assays, further validating *C. elegans* qHTS as a large-scale quantitative profiling method to evaluate *in vivo* compound toxicity.

## Discussion

Here we describe the first example of qHTS employing a multicellular organism. Building on prior work reporting examples using *C. elegans* in chemical library screens (Gosai et al., 2010; Iyer et al., 2019; Knox et al., 2021; Ye et al., 2014), our study attempts to expand the versatility of *C. elegans* as a qHTS-compatible model organism. Towards this end, *E. coli* BGs were employed as a stable, non-replicating nutrient during a multiday screen and LSC provided efficient fluorescent signal detection from *C. elegans* expressing GFP in 384-well microtiter plate format. The resulting dose-response curves enabled more informed prioritization for follow-up chemotype cluster retesting, life-stage, and proteomics analysis. There are certain constraints in applying qHTS to *C. elegans*. For example, the animal number and size distribution described here preclude the surface area/volume limitations of 1536-well plates routinely used in large-scale chemical library qHTS (Inglese et al., 2006). Further, scale-up of synchronized *C. elegans* populations suitable for very large library qHTS would present logistical challenges. However, as demonstrated in this study *C. elegans* qHTS is ideally suited for use with increasingly available high-impact chemical libraries of a moderate size such as approved drugs, protein kinase inhibitors, academic consortium libraries, or environmental chemical collections (Brown et al., 2011; Drewry et al., 2014; Huang et al., 2011; Kearney et al., 2018; Ngan et al., 2019).

LSC was highly effective in scoring worm populations for the construction of DRCs to permit rapid sorting via CC values for subsequent retesting (**Tables S3_S5**). Though not necessary in the current study, additional LSC object parameters, such as aspect ratio and Gaussian distribution of fluorescent signal, can categorize worms based on shape and morphology, e.g., the natural sinusoidal vs. curled phenotype initially observed upon treatment with paralytic agents such as levamisole and albendazole (Lycke et al., 2013; Sant’anna et al., 2013). In addition to phenotypic classifications possible from fluorescently tagged proteins or anatomical regions such as the gut, body wall, phalanx, or neurons, the three fluorescent channel capability of LSC would in principle enable colocalization and multiplexed assay formats (Auld et al., 2006). For example, metabolic incorporation of TAMRA-dATP into genomic DNA or “cell painting” could facilitate a range of phenotypic profiles in *C. elegans* (Gustafsdottir et al., 2013; Schreier et al., 2022). While we conducted the current study with LSC, using automated microscopy as a primary or LSI-coupled output is certainly feasible (Gosai et al., 2010).

In this proof-of-principle study, collectively >1,500 compounds were pharmacologically evaluated for an anthelminthic effect on BG-fed *C. elegans* in qHTS format over a 7-day time course using an LSC readout. From the anti-infectives library, three anti-nematodal agents (**Figure 3**, cpds 1-3 and **Fig. S6A**) were identified through high-quality dose-response CCs thus serving as blinded internal controls. A fourth anti-parasitic compound, praziquantel (cpd 4, **Fig. S7**), administered for trematodes and cestodes infections, though having no reported nematode efficacy (Greenberg, 2005), was not identified as active, further demonstrating assay selectivity.

The chemotype-dependent species selectivity demonstrated in this study highlights the importance of including a structurally diverse selection for each mechanistic drug class to best leverage the power of *C. elegans* as a potential phenolog of human disease in chemical library screening. For example, proteasome and bromodomain (BRD) inhibitors were identified as molecular classes with potential anthelmintic properties. For these, 4 of 11 proteasome and 3 of 7 BRD inhibitors within each class were identified as having high-quality inhibition curves reconfirming in follow-up analysis (**Fig. S6**). Among the 11 proteasome inhibitors in the library, only the four boronic acid-containing chemotypes were active (**Figure 3D**), while the other proteasome inhibitor classes (epoxides, aldehydes, and nitro sulfonic and vinyl amides, **Fig. S10A**) were not identified as active in the primary screen, illuminating the significant species differences reflected in either the mammalian vs. *C. elegans* proteasome sequence or the bioavailability of the chemotype in this organism. For the BRD inhibitors, the differential activity observed among the chemotypes (**Fig. S10B, C**) is not as clearcut as with the boronic acids potentially reflecting various mechanisms including off-target activities.

The primary screening sensitivity of qHTS may be particularly advantageous in assessing drugs and investigational agents in *C. elegans* where potency and efficacy are not optimized for this organism. For example, the monoacylglycerol lipase (MAGL) inhibitor, JZL-184, and nuclear export protein inhibitor analogs, selinexor and verdinexor, were captured by qHTS despite having low potency and efficacy on *C. elegans* viability in the primary 7-point screen (**Table S3**), whereas re-testing in 11-point titration confirms the observation (**Table S5**) which is selective on *C. elegans* over the human cell line (**Fig. S6A**). In the case of JZL-184, whether the effect on *C. elegans* viability is promoted by MAGL inhibition or off-target activity on other serine hydrolases such as FAAH remains to be determined (Long et al., 2009). However, given the dependence of cholesterol trafficking on endocannabinoid metabolism and signaling the observed result is consistent with impaired *C. elegans* development (Baggelaar et al., 2018; Galles et al., 2018).

Not surprisingly, reactive, or promiscuously acting compounds show effects on *C. elegans* viability. Here, molecules including pristimerin, tyrphostin A9, auranofin, and oligomycin A displayed potent and/or efficacious toxicity often, but not always with concomitant activity on the human cell line control (**Fig. S6B, D**), suggesting an application of this assay system in chemical toxicological assessment studies where detection of adverse effects requires a more sophisticated *in vivo* model than possible with cell culture systems.

The pharmacological output of qHTS is well-aligned with subsequent omics-based analysis to aid in the elucidation of the mechanism, illustrated in this study, using MS-based proteomics to interrogate the action of proteasome inhibitors on networks tied to the inhibition of this protein complex and associated cellular processes (**Figure 4E**). The rapid deciphering of a drug or toxic response to an underlying mechanism or potential target protein can substantially help advance the search and development of new therapeutics and potentially flag accompanying liabilities. We anticipate the platform described here will further integrate the use of *C. elegans* orthologous phenotypic models of human disease for pre-clinical drug discovery or repurposing, and in the assessment of potential environmental and human toxins (Golden, 2017; Kim et al., 2018; McGary et al., 2010).

## Methods and Materials

### *C. elegans* culture

*C. elegans* wild-type strain N2 Bristol isolate and strain PE254, feIs4 [*sur-5p*::luciferase::GFP + *rol-6(su1006)*] (Lagido et al., 2009; Lagido et al., 2008; McLaggan et al., 2012), were purchased from the *Caenorhabditis* Genetics Center (CGC; University of Minnesota) and grown according to the Maintenance of *C. elegans* WormBook (Stiernagle, February 11, 2006). Specific details used in this work are elaborated on in the **Supplementary Methods**.

### Compounds and Anti-Infectives Library Preparation

Levamisole, ivermectin and albendazole were purchased from Sigma Aldrich. Stock solutions of all compounds were prepared at 10 mM in DMSO and ivermectin was diluted to a 10 μM working dilution in DMSO. The NCATS Anti-Infectives Collection comprises 643 small molecule compounds having demonstrated activity against viral, bacterial, or eukaryotic pathogens. The library was arrayed as DMSO stock solutions in Echo-qualified 384-well cyclic olefin copolymer plates (Labcyte, Inc.) formatted for qHTS. Specifically, in this case qHTS format was an inter-plate 7-point, 1:5 dilution series spanning high concentrations ranging between 2.5 to 20 mM titrated to 160 nM to 1.28 μM at the low concentration end (Yasgar et al., 2008). “Assay ready” plates were subsequently prepared in 384-well uClear microtiter plates (Greiner BioOne) with compounds acoustically dispensed at 250 nL per well with an Echo 555 in the above inter-plate titration array. Complete details of qHTS plate controls are provided in **Protocol Table S4**. Assay ready plates were sealed and frozen at -80 °C until use.

### Construction of lysis vector and bacterial ghost (BG) production

Lysis plasmid pλP_R_ cI-Elysis was prepared as described previously (Kwon et al., 2005), with some modifications as outlined in **Supplementary Methods** and **Protocol Table S1**. Top10 *E. coli* (Thermo Fisher Scientific, USA) transformed with pλP_R_ cI-Elysis were grown in Luria Broth (LB) (Thermo Fisher Scientific, USA) supplemented with ampicillin (50 μg/mL) at 28 °C with 200 rpm shaking until OD_600_=0.4-0.6 at which point temperature was shifted to 42°C for ∼2 h to induce the Lysis E gene product. Further details including quality control and storage conditions are described in **Supplementary Methods** and **Protocol Table S2**.

### Calibration of COPAS biosorter for *C. elegans*

N2 *C. elegans* eggs were axenized from a mixed population of worms grown on ten NGM plates according to the Maintenance of *C. elegans* WormBook (Stiernagle, February 11, 2006). Eggs were hatched overnight in 9 mL M9 buffer at 20°C with 65 rpm shaking then collected with centrifugation at 1000 rpm for 5 min. Worms were resuspended in 100 mL S medium with 250 mg frozen pellet of OP50. Worms were grown at 20°C with vigorous shaking and 20 mL of worms collected at respective time points. Early-L1s were collected at 6 h, mid-L2s collected at 20 h, mid-L3s collected at 28 h, mid-L4s collected at 38 h and adults collected at 48 h (Sulston, 1988). Each worm harvest was loaded on a COPAS Flow Pilot (FP-500) biosorter (Union Biometrica, Inc., MA) in 50 mL tubes and a bulk sort of 1000-5000 objects was performed. Gating was manually set within the experimental protocol file to collect each individual life stage based on Time-of-Flight (TOF) and Extinction Coefficient (Extinction). Full details are provided in **Supplementary Methods**.

### Characterization of *E. coli* BGs as a *C. elegans* nutrient source

NGM plates were treated with 200 μL of *E. coli* BGs or OP50 live bacteria incubated ±DiBAC4(3) in PBS buffer and N2 *C. elegans* were expanded on the respective NGM plates with sterile chunking so that each plate received 40-60 worms per condition. After five days of growth worms were collected from respective plates, washed, and resuspended in 10 mL 1X S basal buffer for COPAS biosorter analysis and imaging on a GE InCell analyzer 2200 (GE Healthcare). See **Supplementary Methods** for full details.

### Assay development employing LSC and BG nutrient source

A mixed population of *C. elegans* strain PE254 was dispensed by life stage at 10 GFP-expressing worms per well into respective wells of 384-well uClear microtiter plates with the COPAS biosorter. *E. coli* BGs or OP50 live bacteria in S medium were added at designated concentrations to respective plates at 30 μL per well with a Capp pipette (BioVentures, Inc.). Levamisole, ivermectin or albendazole titrations were transferred (300 nL) to corresponding microplate columns with a Mosquito HTS nanoliter liquid handler (STP Labtech). DMSO and 50 μM final concentration levamisole were added to designated columns and rows as vehicle and toxicity controls, respectively as detailed in **Protocol Table S3**. Plates were incubated for seven days at 21°C and 65 rpm in an Incu-Shaker (Benchmark Scientific) and imaged every 24 – 72 h on an Acumen eX3 laser scanning cytometer (STP Labtech) using settings outlined in **Supplementary Methods**.

### Primary qHTS

For each library collection, *C. elegans* strain PE254 were collected from twelve NGM plates as above, transferred to 250 mL S medium supplemented with OP50 *E. coli* pellet collected from 2 L overnight culture, and grown for five days at 21°C with vigorous shaking. Eggs were axenized as above and hatched overnight in M9 buffer at 21°C and shaken at 65 rpm. Assay ready plates were thawed and allowed to equilibrate to ambient temperature. 30 μL/well BGs in S medium at an OD_600_=0.85 were manually added across plates with a Capp pipette. Synchronized L1 worms from above were collected, diluted, and plated at 10 worms/30 μl/well with Capp pipette into assay ready microtiter plates pre-plated with BGs and compounds (240-fold dilution). Plates were read on an Acumen eX3 LSC as above with resolution settings of X=2 μm and Y=16 μm within 4 h of plating and at 3- and 7-day time points (see **Protocol Table S4** for further detail). The Anti-Infectives library was tested as a 7-point, 1:5 inter-plate titration and the Tox21 sub-library as a 7-point, 1:5 intra-plate titration.

Mammalian cell toxicity was evaluated in HEK293 cells (ATCC) using ATP levels as an indicator of metabolically active cells. Assays were performed in 1536-well plates (Greiner white/solid bottom high base) and luminescence measurements collected on a ViewLux (Perkin Elmer). See **Protocol Table S5** for details.

### Data analysis for assay development and qHTS

Assay statistics including signal-to-background, signal-to-noise, and Z’ (Southall et al., 2009; Zhang et al., 1999) were calculated for each life stage of *C. elegans* strain PE254 grown on various bacteria media treated with the intra-plate control compounds, 50 μM levamisole toxicity control and DMSO neutral control. Concentration-response curves for average total filtered FL-2 object area (“GFP area”) and total filtered FL-2 number of objects (“number of worms”) for each respective compound were fit with the nonlinear regression log (agonist) vs. response – variable slope (four parameters) algorithm in Prism (GraphPad Software).

Data from the compound libraries screens were normalized by plate to corresponding intra-plate controls as previously described (Inglese et al., 2006). Assay statistics were calculated using the same respective controls. Curve fits assignments of -1.1, -1.2, -2.1, -2.2 and -3 were considered active and visually confirmed. These data were refit in Prism with nonlinear regression log(agonist) vs. response – variable slope (four parameters) fit. See **Supplementary Methods** for further detail.

### Follow-up confirmation testing and *C. elegans* life-stage analysis

*C. elegans* strain PE254 were collected from six NGM plates, washed, and eggs were axenized as above and hatched overnight in M9 buffer at 21 °C and shaken at 65 rpm. Synchronized L1 worms and BGs were added to assay ready plates prepared as above with an 11-pt, 1:3 intra-plate titration of 96 compounds identified as active from primary qHTS and plates and read on an Acumen eX3 LSC as above within 1 h of plating and every 24 h for a 7-day time course (**Protocol Table S6**).

For life-stage analysis, after a 7-day compound treatment in 6-well culture plates seeded with a mixed population of 2,000 *C. elegans* strain PE254, life-stage distributions were determined by Time-of-Flight (TOF) and Extinction Coefficient (EC) using the COPAS FP BioSorter, followed by bulk collection in a 50 mL conical tube for use in proteomics studies. See **Protocol Table S7** for further details.

### Proteomics sample preparation

Compound and control treatment groups comprised of three biological replicates each were prepared for proteomic digestion by trypsin prior to HPLC-MS/MS analysis. The groups consisted of untreated and DMSO control groups and a bortezomib (4 µM), trametinib (1 µM), LY411575 (500 nM), selumetinib (10 µM) and selinexor (20 µM) treated groups. NP40 cell lysis buffer (Thermo Fisher Scientific) with protease inhibitor cocktail (Sigma) was added to each worm sample for LN2 manual grinding protein extraction and quantification with BCA. Approximately 30 µg of each lysate was reduced, alkylated, acetonitrile precipitated, and trypsinized as further described in **Supplementary Methods**.

### HPLC-MS/MS analysis

All proteomic HPLC-MS/MS analysis was performed using an UltiMate 3000 RSLCnano LC system coupled to the Orbitrap Fusion Lumos Tribrid Mass Spectrometer equipped with the Nanospray Flex ion source (Thermo Fisher Scientific, Waltham, MA, U.S.A.) as previously described (Burns et al., 2021) and detailed in **Supplementary Methods**.

### Data Analysis for proteomics

Proteome Discoverer software suite (v2.4, Thermo Fisher Scientific) with Sequest algorithm were used for peptide identification and quantitation as previously described and detailed in **Supplementary Methods** (Burns et al., 2021). The HPLC-MS/MS raw data files and associated search results have been deposited to the ProteomeXchange Consortium via the PRIDE partner repository with the dataset identifier PXD033552 (Perez-Riverol et al., 2019).

## Supporting Information

This material is available free of charge via the Internet at http://

- Characterization of *C. elegans* growth on *E. coli* ghosts, laser scanning cytometry detection parameter analysis, mammalian cell lines cytotoxicity qHTS, Re-test summary statistics and concentration–response curves, (PDF).
- Additional details of assay development statistics, primary qHTS summary table, follow-up data tables, and assay protocol tables (XLSX).

### Accession Codes

The qHTS data reported in this study has been deposited to the PubChem (https://pubchem.ncbi.nlm.nih.gov/) database with the identifier AIDs 1745854 and 1745855. The HPLC-MS raw data files and associated search results have been deposited to the ProteomeXchange Consortium via the PRIDE partner repository (http://proteomecentral.proteomexchange.org/cgi/GetDataset) with the dataset identifier PXD033552. The pλPR cI-Elysis plasmid has been deposited in Addgene under accession code 176891.

## Acknowledgements

This research was supported by the Intramural Research Program of the National Center for Advancing Translational Sciences (NCATS), National Institutes of Health (NIH) under project 1ZIATR000052 (J.I.). *C. elegans* strains were obtained from the *Caenorhabditis* Genetics Center (CGC), funded in part by the NIH Office of Research Infrastructure Programs (P40 OD010440). We thank NCATS Compound Management for assay-ready library plates, C. Tanega for PubChem depositions, D. Leja (NHGRI) for qHTS illustration, and Drs. J.L. Dahlin (NCATS), C.P. Venditti (NHGRI) and A. Golden (NINDS) for their comments on the manuscript. Graphic abstract and Figure 1C were created with BioRender.com.

## Footnotes

1 ∼4,500 proteins listed in the UniProt Knowledgebase (UniProtKB)

